# Directional Immobilisation of SpyTag Bacteriophage on PDMS surfaces for Phage based Microfluidics

**DOI:** 10.1101/2024.10.11.617866

**Authors:** Sahan B.W. Liyanagedera, Joshua Williams, Joseph P. Wheatley, Alona Yu Biketova, Antonia P. Sagona, Tamas Feher, Vishwesh Kulkarn

**Affiliations:** School of Biological Sciences, University of Edinburgh, United Kingdom; School of Engineering, University of Warwick, United Kingdom; School of Life Sciences, University of Warwick, United Kingdom; Jodrell Laboratory, Royal Botanic Gardens, United Kingdom; HUN-REN Biological Research Centre, Szeged, Hungary; School of Engineering, Kings College London, United Kingdom

**Keywords:** Microfluidics, PDMS, Biosensor, Point of Care, SpyTag-SpyCatcher, CRISPR/Cas, Engineered Bacteriophage, Phage Therapy, Immobilisation

## Abstract

The increasing incidence of bacterial infections caused by antibiotic-resistant pathogens worldwide underlines the need to develop novel diagnostic tools enabling the early initiation of targeted antimicrobial therapy. One promising possibility is to unite the high specificity and sensitivity of phage-based applications with the speed and sensitivity provided by microfluidic devices. As a prerequisite of developing such systems, we aimed at the directional immobilization of phages on the surface of Polydimethylsiloxane (PDMS), a material commonly used for building such devices. Our work utilised the covalent interaction between two proteins: SpyTag, genetically encoded on the capsid of the phage, and BslA-SpyCatcher fusion protein, purified and surface displayed on PDMS. We demonstrate a simple methodology for the directional tail up immobilisation of SpyTagged Phage on to user defined locations on the surface of a PDMS device and subsequent on chip capture and infection of a cognate host. Our technique serves to illustrate a generally applicable solution to develop the next generation of phage based bio-sensors.

## 1 Introduction

Pathogenic bacterial infections require early and rapid detection to maximise the chances of therapeutic success and survival. However, conventional bacterial detection methods are highly complex, require trained personnel and are time-consuming.(*1*) While advanced molecular biology techniques such as polymerase chain reaction (PCR) and enzyme-linked immunosorbent assay (ELISA) have significantly reduced detection times and increased sensitivity, these approaches still suffer from high cost and slow turnaround from sample collection to confirmatory result. Therefore, the development of rapid, sensitive, cost-effective, and highly specific bacterial detection systems is of high importance in clinical diagnoses, food industries and bio-counter terrorism.(*2*)

Bacteriophages (phages) are viruses that infect bacteria.(*3*) With relatively narrow host ranges, phages could be used to detect the presence of specific bacterial pathogens if they could be integrated into a low-cost biosensor. Biosensors usually consist of a biological recognition component, a transducer and an electrical system which amplifies, processes, and computes the signal.(*1*) The use of a bacteriophage as a detection probe or recognition component has several advantages for rapid bacterial screening including (1) extreme specificity to cognate host, (2) massive increase in progeny phage (thus a detectable signal) from a single infection event, (3) high tolerance to harsh environmental conditions and processing, and (4) simple and inexpensive large-scale production.(*1*) These advantages highlight and underpin the use of bacteriophages in the development of biosensors for bacterial detection. While there have been a number of studies that have capitalised on the use of bacteriophages as biosensors.(*2, 4 –10*), the current state-of-the-art utilising such methods still does not meet the sensitivities required for medical applications, nor do they demonstrate the speed and simplicity of use needed in a point of care (P.O.C) diagnostic device. The fabrication of such a device require the development of methods for the immobilisation of bacteriophages which can be achieved through chemical, physochemical or electrostatic mechanisms between the virus and the surface in question.(*11*) Nonetheless, such methods do not necessitate directional immobilisation, whereby the bacteriophage is oriented in a manner so as to maximise contact between its receptor components and their target host. This is especially true in the case of tailed phages, which comprise nearly 96% of all phages described to date, for which simple and orthogonal strategies for directional immobilisation onto materials of interest are highly sought after.(*3*)

In order to address these limitations, we demonstrate for the first time a method for the directional immobilisation of phage on Polydimethylsiloxane (PDMS) surfaces, a material commonly used to fabricate microfluidic based diagnostic devices. This foundational technique is a critical step toward the development of phage immobilised microfluidic P.O.C platforms which could serve as an end-user device for simple, rapid and highly sensitive detection of pathogenic bacteria.

As a proof of concept, we focus on the directional immobilisation of bacteriophage K1F on PDMS, for the on-chip capture and detection of *E. coli* K1, a Gram-negative pathogen responsible for a wide range of diseases, including sepsis, urinary tract infections, inflammatory bowl syndrome and neonatal meningitis.(*12 – 15*) Thus the rapid diagnosis and initiation of treatment in the form of classical antibiotics or phage therapy are critical to survival and favourable prognosis. Furthermore, the recent prevalence of clonal variants and antibiotic resistance has generated a need to develop new methods to quickly identify the specific pathogenic strain causing infection and provide targeted antimicrobial treatment.(*11*)

In our approach, we immobilise K1F phage onto the surface of PDMS chips, using a well-characterised system for protein immobilisation; the SpyCatcher/SpyTag System (Figure 1).(*16*) For this purpose, we utilised an engineered variant of K1F, namely K1F-STG, whereby GFP and SpyTag were integrated onto the capsid head of phage K1F. (*17*) Through a combination of rational protein design for directional immobilisation onto PDMS and a simple fabrication methodology for user defined deposition of immobilised phage; we aim to provide generally applicable solutions to increase sensitivity and reduce the time of detection for the next generation of phage-based biosensors.

**Figure 1:**
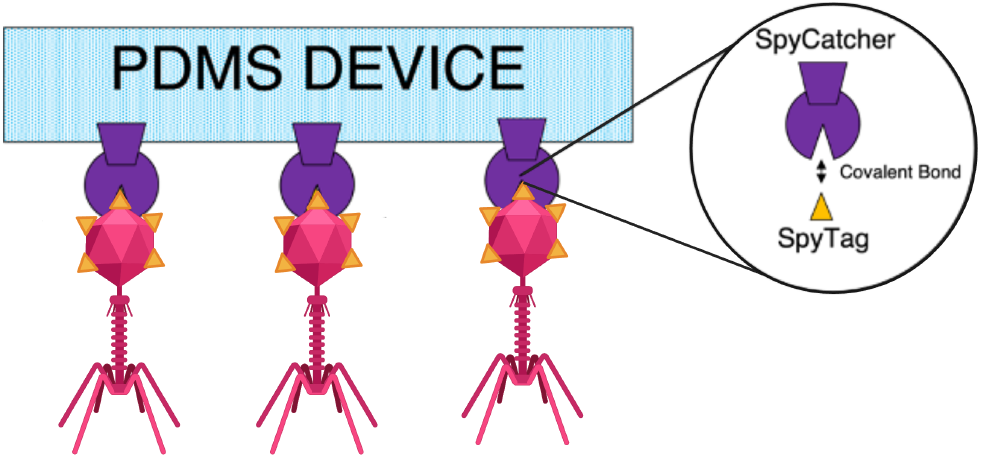
A general strategy for directional immobilisation of Phage on PDMS surfaces. The creation of a phage based point of care device for bacterial detection whereby surface displayed SpyCatcher fusions enable the covalent coupling of SpyTagged phage at user defined locations.

## 2 Results and discussion

A principle challenge in building a phage based diagnostic device stems from the difficulty in ensuring directional immobilisation of phages on the material of interest. Directional immobilisation improves stability whilst maximising the contact of bacteriophage receptor components to their target host. Nonetheless, to the best of our knowledge, no results are available for the immobilisation of bacteriophages on to PDMS, a material commonly used in the preparation of microfluidic diagnostic devices. The results presented herein constitute an important preliminary step in this direction. The core strategy of our work was to rely on the highly specific and covalent SpyTag-SpyCatcher interaction to immobilize bacteriophages on solid PDMS surfaces. This required two innovations: the expression of SpyTag protein on the virion head, as well as developing an optimal fusion protein of SpyCatcher to facilitate its oriented immobilization in a surface-exposed manner on PDMS, thereby enabling effective covalent bonding with SpyTagged phages at user defined locations.

### Engineering SpyTagged K1F Phage

To enable directional (tail-up) immobilisation of K1F phage, we decided to incorporate the SpyTag gene into the phage capsid head through homologous recombination and subsequent CRISPR/Cas9 selection.(*17*) The SpyTag protein is one half of the powerful protein conjugation pair termed the Spy-Tag/SpyCatcher system.(*16*) The system is derived by splitting the *Streptococcus pyogenes* fibro-nectin-binding protein FbaB into two functional domains, viz., SpyTag and Spy-Catcher, that form a spontaneous isopeptide bond between Lysine and Asparagine residues.

We previously demonstrated that this ultrastrong covalent interaction is an ideal method for capsid decoration of SpyTagged K1F phage to generate targeted phage therapies.(*17*) We therefore postulated that the same mechanism could be utilised for facile directional immobilisation of the SpyTagged phage on to surfaces containing SpyCatcher fusion proteins. In this instance, the fusion is targeted to the non essential minor capsid protein (gene10b), which is formed by a programmed -1 translational frameshift in the 3’ region of g10, adding a further 44 amino acids to the protein sequence. The minor capsid protein is found in lower abundance than its major counterpart, assembling at the apexes of the capsid head. While the incorporation of SpyTag to the major capsid protein would provide significantly more attachment points for covalent interactions, our previous work demonstrated that this fusion is unstable and is lost following a few rounds of propagation.(*17*) As such, we chose the K1F-STG variant, whereby GFP and SpyTag is incorporated into the minor capsid protein via homologous recombination and CRISPR/Cas9 mediated selection as previously described.(*17*) .

The GFP incorporated in the capsid of K1F-STG serves as a simple biosensor whereby infection of target bacteria on the PDMS chip by immobilised K1F-STG will result in a easily detectable fluorescent signal. Furthermore, we incorporated a Tobacco Etch Virus (TEV) protease target site immediately before the SpyTag gene. This strategic design allows for the proteolytic cleavage and subsequent removal of any covalently bonded SpyCatcher-SpyTag fusion from the capsid head. This feature provides a controlled release mechanism for immobilized phage particles, which could be essential for applications such as on-chip cell-free phage synthesis and purification.(*17, 18*) Donor cassette sequence and design for creation of K1F-STG are provided in Supporting Information Table S3 and Figure S1 respectively.

### PDMS Surface Decoration with Spy-Catcher Fusion Proteins

Following the creation of K1F-STG phage incorporating GFP and SpyTag motifs on their capsid heads, we focused on the development of a simple strategy for the immobilisation of SpyCatcher proteins on the surface of PDMS chips. During the development of an immobilisation strategy, inspiration was drawn from the seminal work by Blum and co-workers, which involves the direct transfer of dried protein/salt solutions to the PDMS interface during polymer curing.(*19*) This simple and versatile method utilises a hydrophobic array template or cast on to which the protein of interest is spotted at user-defined locations. PDMS is subsequently poured on top of the dried protein spots and allowed to set. The cured PDMS is then peeled away, revealing immobilised proteins at the polymer surface (Figure 2).

**Figure 2:**
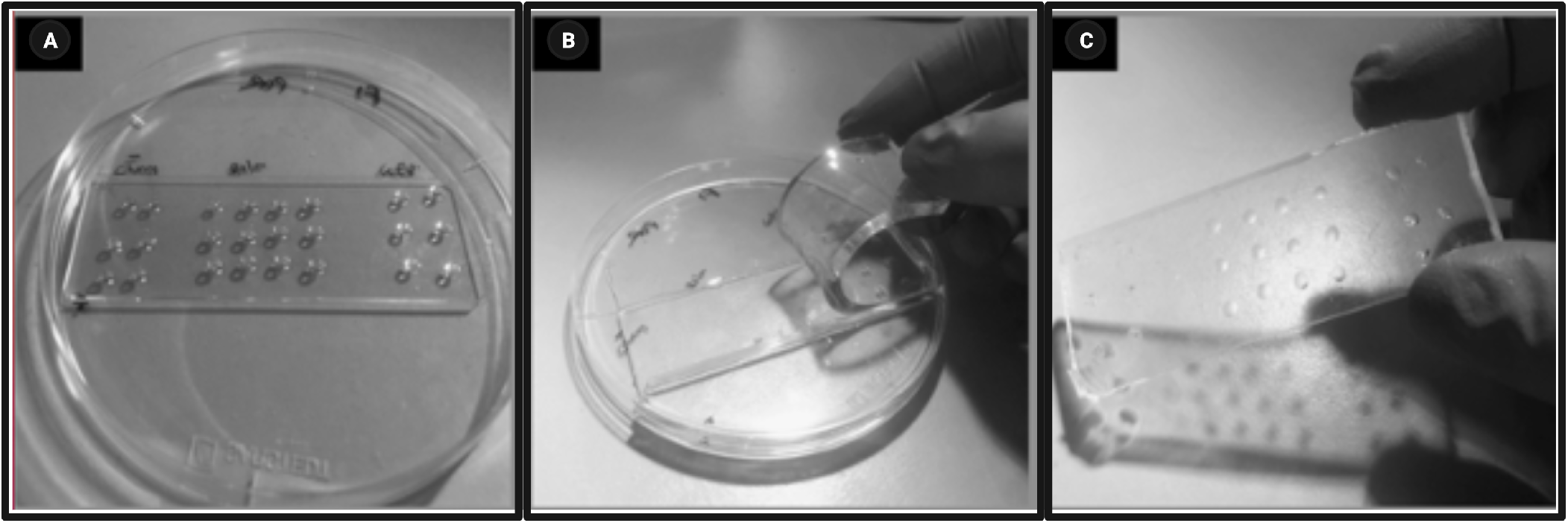
Simple strategy for the immobilization of SpyCatcher proteins on PDMS. SpyCatcher proteins are spotted onto a hydrophobically treated microscope slide before pouring over PDMS **(A)**. Once cured, PDMS is cut around the slide and peeled back carefully. This process results in the capture and surface display of SpyCatcher proteins **(B)**. Cured PDMS device with surface immobilised SpyCatcher proteins **(C)**.

We initially used a SpyCatcher-mCherry protein spotting solution to replicate this method using a standard single-channel silicone microfluidic device mould that was treated with a rain-repellent hydrophobic agent. Following the drying of the SpyCatcher-mCherry protein solution on the silicon wafer mould, PDMS was poured over and allowed to cure as previously described. Following curing, the devices were peeled off the mould and cut into squares ready for surface capture experiments with SpyTag fusions.

As a simple test to check for surface display of the SpyCatcher-mCherry protein, we incubated a solution of purified GFP-SpyTag proteins at the sites of SpyCatcher-mCherry surface immobilisation, to enable covalent linkage to occur between SpyCatcher and SpyTag (Supplementary Figure S4). These devices were then thoroughly washed in 2L of sterile water and sonicated for 30 minutes to remove any GFP-SpyTag proteins that had not covalently bound to SpyCatcher-mCherry and retention of GFP fluorescence was observed using a protein gel-imager.

These experiments led to little to no capture of SpyTag-GFP on the PDMS surface as was observed via low GFP fluorescence (Figure 4B). It was hypothesised that the poor retention of GFP-SpyTag was due to low levels of SpyCatcher-mCherry being surface displayed during the PDMS curing process. Furthermore, the initial study by Blum and coworkers identified both a moulding effect and a chemical mechanism by which the proteins in the spotting solution are captured by the PDMS during the curing process.(*19*) The latter chemical mechanism was determined to occur via the covalent linkage of polymer to the lysine residues on the protein, through the poisoning of the Kardstedt catalyst during polymer curing.(*19*) It is therefore likely that during the entrapment of SpyCatcher-mCherry onto the PDMS surface, the lysine in the active site of SpyCatcher is partaking in the covalent coupling of Spycatcher-mCherry to PDMS. In such an instance, while SpyCatcher-mCherry may indeed become entrapped at the surface of the PDMS, the available “active” SpyCatcher-mCherry proteins with free lysine residues for covalent coupling to SpyTag, are drastically reduced.

In order to circumvent this problem and decrease the likelihood of the active site of SpyCatcher being involved in the poisoning of the Kardstedt catalyst during PDMS curing, SpyCatcher was fused to the bacterial hydrophobin BslA, a protein that readily forms monolayers on hydrophobic surfaces.(*20*) It was envisioned that by fusing SpyCatcher to the hydrophobic terminus of BslA and spotting this protein solution on to a hydrophobic surface, the dried protein monolayer will by virtue of rational protein design, orient the SpyCatcher protein away from the PDMS that is poured on top. In this manner, the orientation of the SpyCatcher protein will be fixed within the dried monolayer and increase the chances of active SpyCatcher proteins being surface displayed on the PDMS chip (Figure 3).

**Figure 3:**
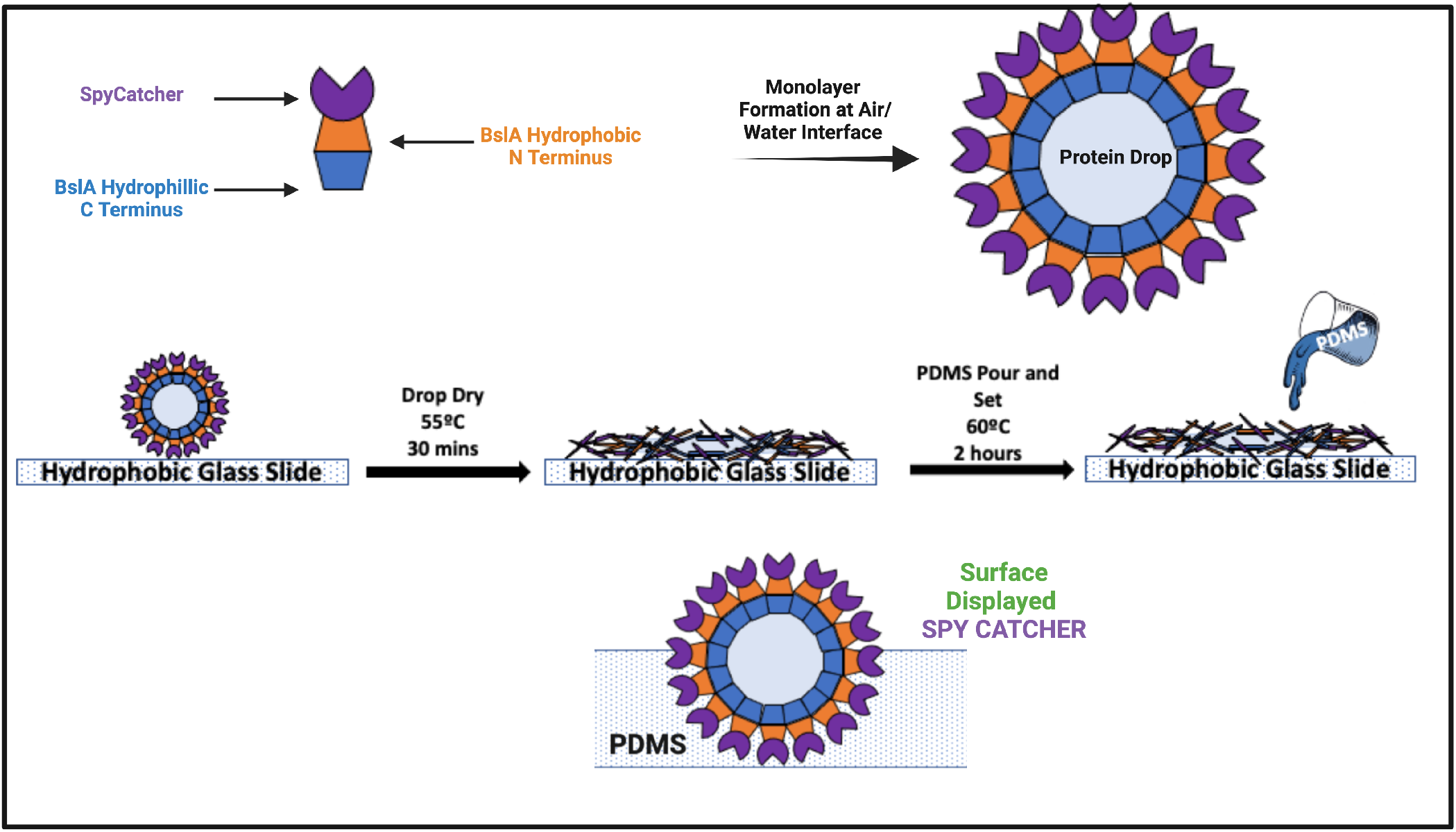
SpyCatcher-BslA rational protein Design for directional immobilisation. The fusion of SpyCatcher to the hydrophobic terminus of the BslA protein **(Top)** ensures the correct orientation of the SpyCatcher protein during PDMS curing **(Centre)**. In this way the reactive groups of SpyCatcher are oriented outwards and surface-displayed and thereby available to react with SpyTagged proteins **(Bottom)**. The orientation of SpyCatcher in this manner also prevents the lysine of the SpyCatcher from partaking in covalent coupling with the PDMS polymer.

Indeed, this strategy led to a significant improvement in the retention of GFP-SpyTag by way of covalent linkage to surface-displayed SpyCatcher-BslA on defined locations on PDMS surfaces (Figure 4). Interestingly, while Blum et.al describes an increase in protein retention on the surface of the PDMS at higher spotting solution concentrations, we found that in the case of the immobilisation of SpyCatcher, the orientation of the protein during curing has a far greater impact on surface display than the protein concentration used. This is clearly illustrated by the fact that while the spotting solutions of SpyCatcher-BslA had a concentration of 1.5 mg/mL, it far outperformed spotting solutions of SpyCatcher-mCherry with concentrations as high as 40 mg/mL.

**Figure 4:**
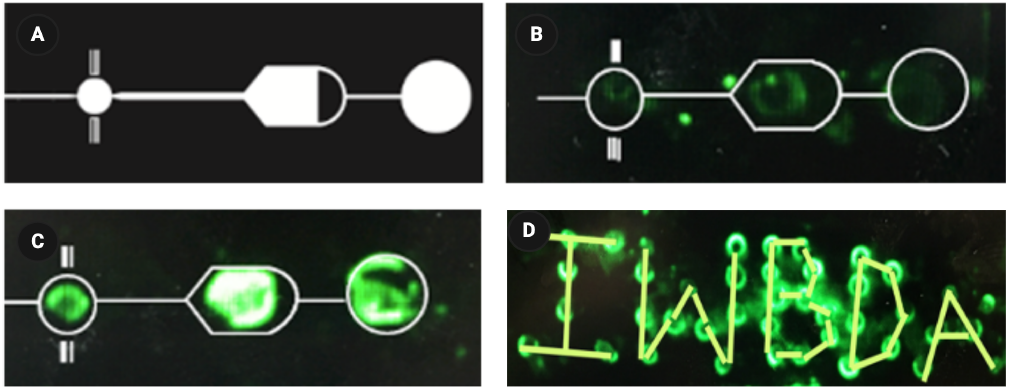
SpyCatcher-BslA surface display enables capture of GFP-SpyTag on PDMS surface. Design of a silicon wafer microfluidic device mould with three chambers for fusion protein immobilisation **(A)**. Low retention of GFP-SpyTag on a PDMS device using 40 mg/mL SpyCatcher-mCherry spotting solution was attributed to potential covalent interactions of fusion with PDMS during curing **(B)**. Enhanced retention of GFP-SpyTag using SpyCatcher-BslA spotting solution **(C)**. Demonstration of location-specific immobilisation on PDMS, spelling “IWBDA” with surface captured GFP-SpyTag **(D)**.

### SpyTag Phage Immobilisation on PDMS surface

Following the establishment of the immobilisation and surface exposure of SpyCatcher-BslA proteins, the method was subsequently used for the directional immobilisation of K1F-STG onto the surface of PDMS devices. For these experiments, a custom mould was created by gluing together 10 glass cover-lips stacked on top of each other onto the centre of a petri dish (Figure 5).

**Figure 5:**
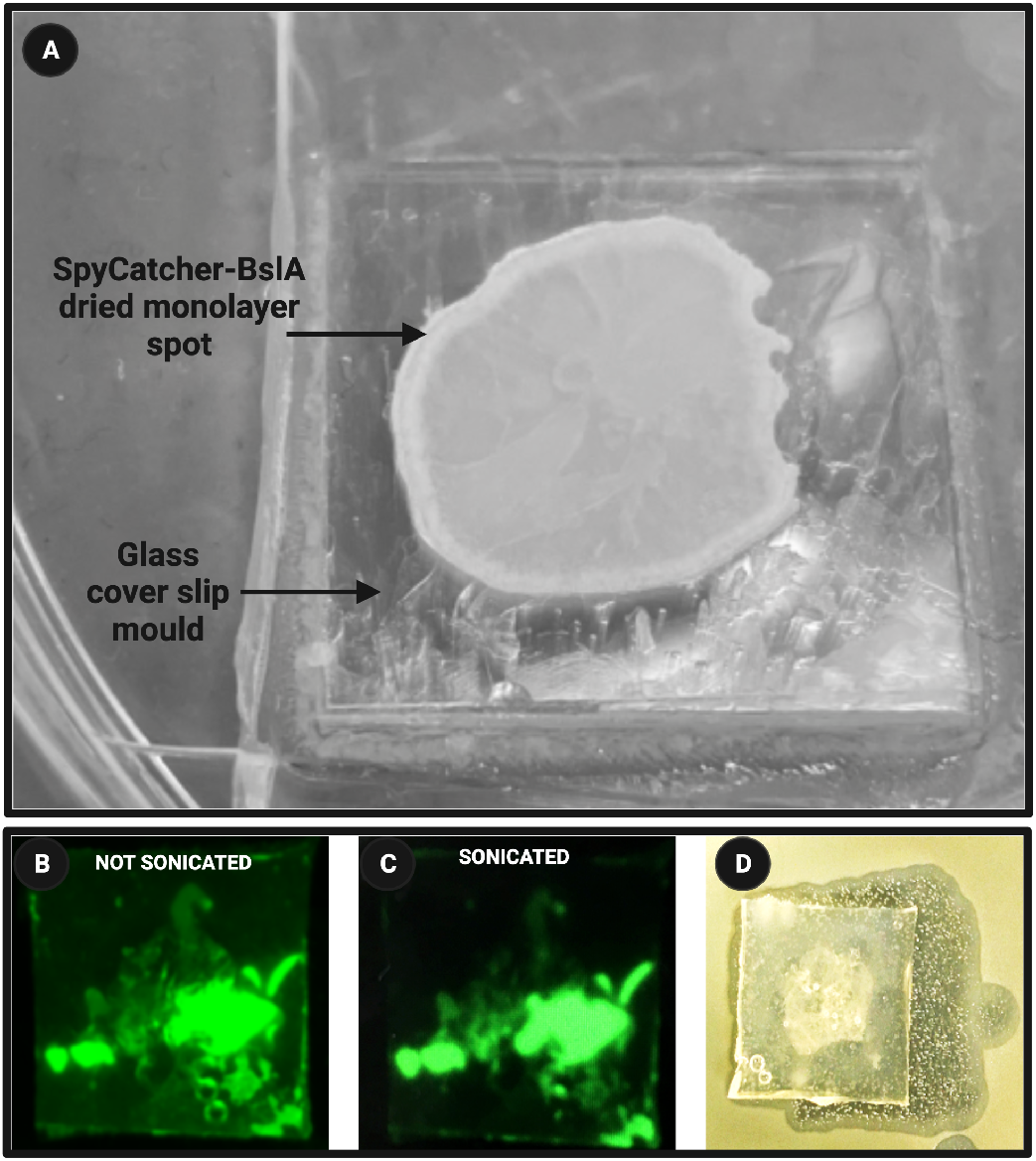
Macroscopic visualisation of directionally immobilised K1F-STG. SpyCatcher-BslA protein in carbonate buffer dried on a custom mold, forming a monolayer with the Marangoni effect causing protein accumulation at the droplet’s periphery clearly visible **(A)**. Images captured using a protein gel imager show green fluorescence on the PDMS surface before **(B)** and after **(C)** sonication. Captured fluorescence includes autofluorescence from SpyCatcher-BslA and fluorescence from K1F-STG. Despite the harsh wash and sonication steps, K1F-STG remains immobilised on the PDMS surface as demonstrated by the ability of the chip to clear lawns of EV36 bacteria **(D)**. These results indicate the strength of the covalent interaction between the SpyTag on the capsid of the phage head and the surface immobilised SpyCatcher proteins.

The protein spotting solution was dried on top of the coverslip mould and PDMS devices were prepared as previously described. Next solutions of PEG purified K1F-STG at 9 Log PFU/mL were incubated on the surface of the device containing immobilised SpyCatcher-BslA proteins. In order to assess the strength of the interaction between K1F-STG and surface-exposed SpyCatcher, chips prepared in this manner were washed in 2L of water and sonicated in a sonicating water bath for 30 minutes. During this step, any passively absorbed K1F-STG phage, as well as K1F-STG phage forming non-specific interactions with the treated surface of the PDMS, should be removed. Indeed, a slight reduction in the surface area of the PDMS chip containing immobilised SpyCatcher-BslA protein was observed at the macroscopic scale in a gel dock imager (Figure 5B and C). Nonetheless, the remaining immobilised phage remained active on the device even after the harsh processing steps and was able to clear the lawns of its host on agar plates (Figure 5D)

Despite the apparent success at the macroscopic level in the directional immobilisation of K1F-STG on PDMS surfaces decorated with SpyCatcher-BslA, fluorescence microscopy of these devices revealed little immobilised phage on the material surface (Supplementary Figure S5). This low capture efficiency was attributed to an unexpected phenomenon that arose due to the drying behaviour of the protein droplets during device fabrication. It was observed that the SpyCatcher-BslA protein solutions dried on the mould were subject to the Marangoni or “coffee ring” effect, where proteins predominantly accumulated at the edges of the droplet. This phenomenon is commonly seen in evaporating liquid droplets containing suspended particles and results from a radical capillary flow that transports material to the periphery as the droplet evaporates.(*21*) The Marangoni effect is clearly visible at the macroscopic scale when looking at a dried SpyCatcher-BslA protein spot (5A).

Furthermore, when the cured PDMS was feeled off the mould, it was observed that the dried SpyCatcher-BslA monolayer ruptures and only a portion of the monolayer gets captured by the PDMS during curing. As such, the SpyCatcher-BslA proteins were primarily retained only at the outer rings of the captured droplet (Figure 5A). This partial capture of the dried protein droplet thus significantly reduced the amount of surface exposed SpyCatcher available for subsequent interactions with SpyTagged K1F phage, thereby limiting the immobilisation capacity of the device.

Taking into consideration these observations, it was postulated that changing the fusion orientation to BslA-SpyCatcher would result in the greatest degree of reactive SpyCatcher moieties on the surface of the PDMS (Figure 6). Indeed, despite the persistence of the coffee ring effect during fabrication with this new fusion orientation, the alteration enhanced exposure of SpyCatcher molecules on the surface of PDMS, improving their accessibility for binding and directional immobilisation of K1F-STG as indicated by the presence of sharp punctate green signals in multiple positions on all devices tested. The BslA-SpyCatcher orientation thus far outperforms the SpyCatcher-BslA orientation for surface display of SpyCatcher as indicated by the substatial increase in GFP fluorescence from covalently linked K1F-STG on devices made using BslA-SpyCatcher (Compare Supplementary Figure S5 and Figure 7).

**Figure 6:**
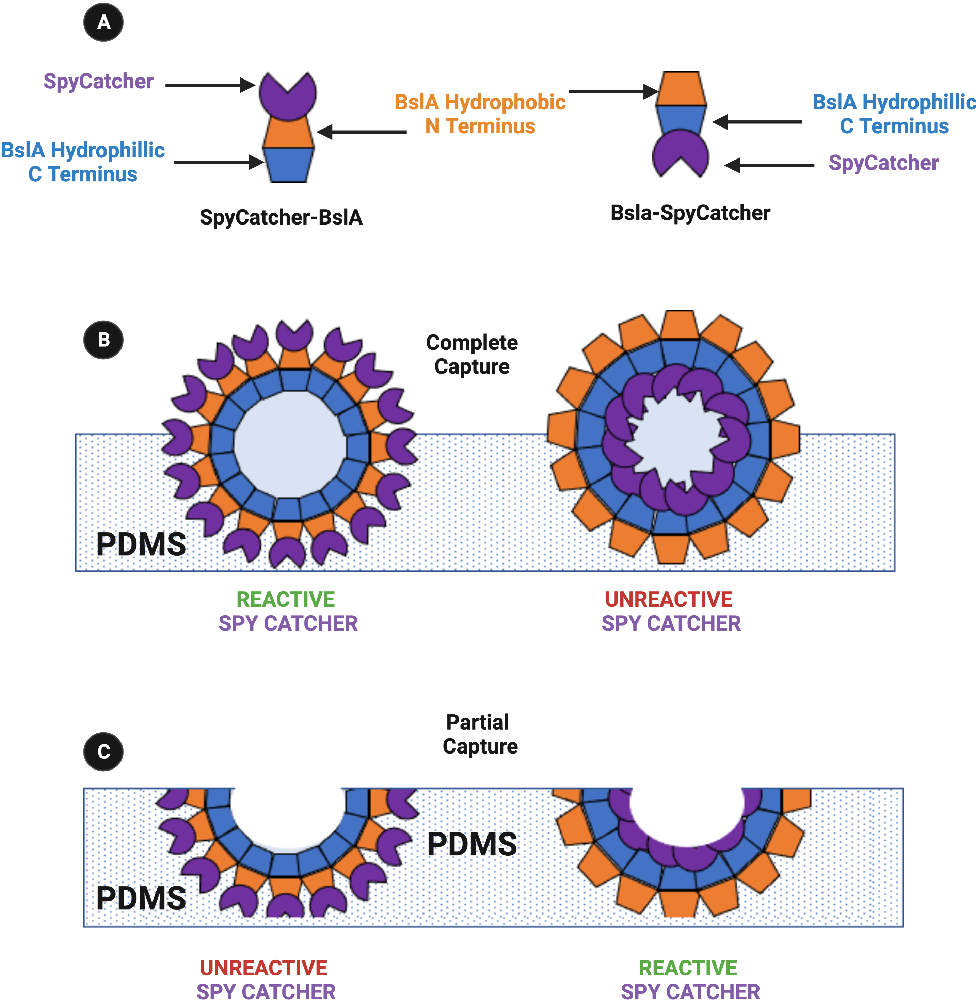
Predicted orientations of PDMS captured Spycatcher fusions to BslA. Spy-Catcher was fused to either the hydrophobic or hydrophilic terminus of BslA to create SpyCatcher-BslA and BslA-SpyCatcher respectively **(A)**. The drying of these protein solutions on a hydrophobic surface and subsequent complete capture by PDMS would result in more surface exposed Spycatcher when using SpyCatcher-BslA compared to BslA-SpyCatcher **(B)**. However, during method establishment, it was observed that the BslA monolayer ruptures and only a portion of the monolayer gets captured by the PDMS during curing. In such an instance, it is expected that the BslA-SpyCatcher fusion orientation would result in the greatest degree of reactive SpyCatcher moieties on the surface of PDMS **(C)**.

**Figure 7:**
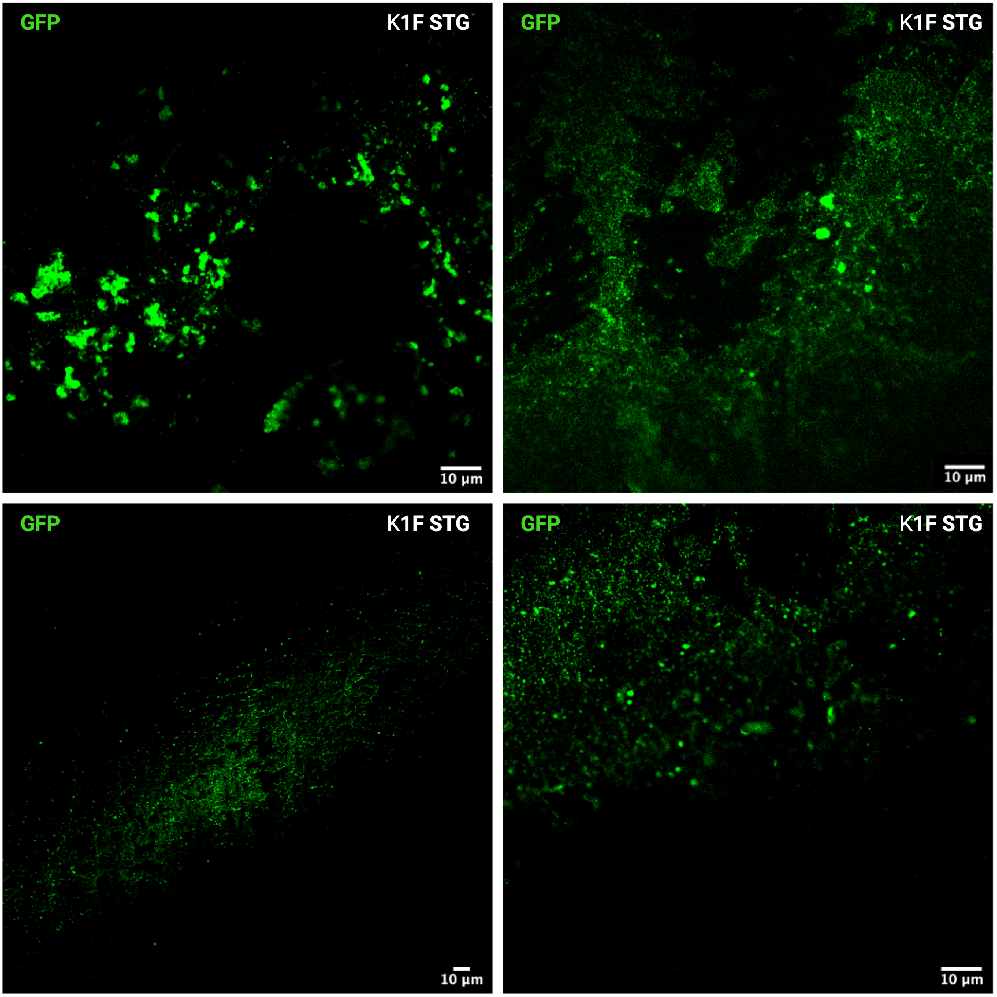
Enhanced capture of K1F-STG by PDMS surface immobilised BslA-SpyCatcher. Representative fluorescence images taken at two different locations on the surfaces of two different PDMS devices made with BslA-SpyCatcher spotting solution. The use of BslA-SpyCatcher as the capture protein in the creation of PDMS devices led to substantial improvements in the surface display of SpyCatcher as indicated by the substantial increase in GFP fluorescence from covlaently linked K1F-STG compared to when SpyCatcher-BslA fusion orientation was used. The bottom left panel was captured at x40 magnification. All other panels were captured at x100 magnification

**Figure 8:**
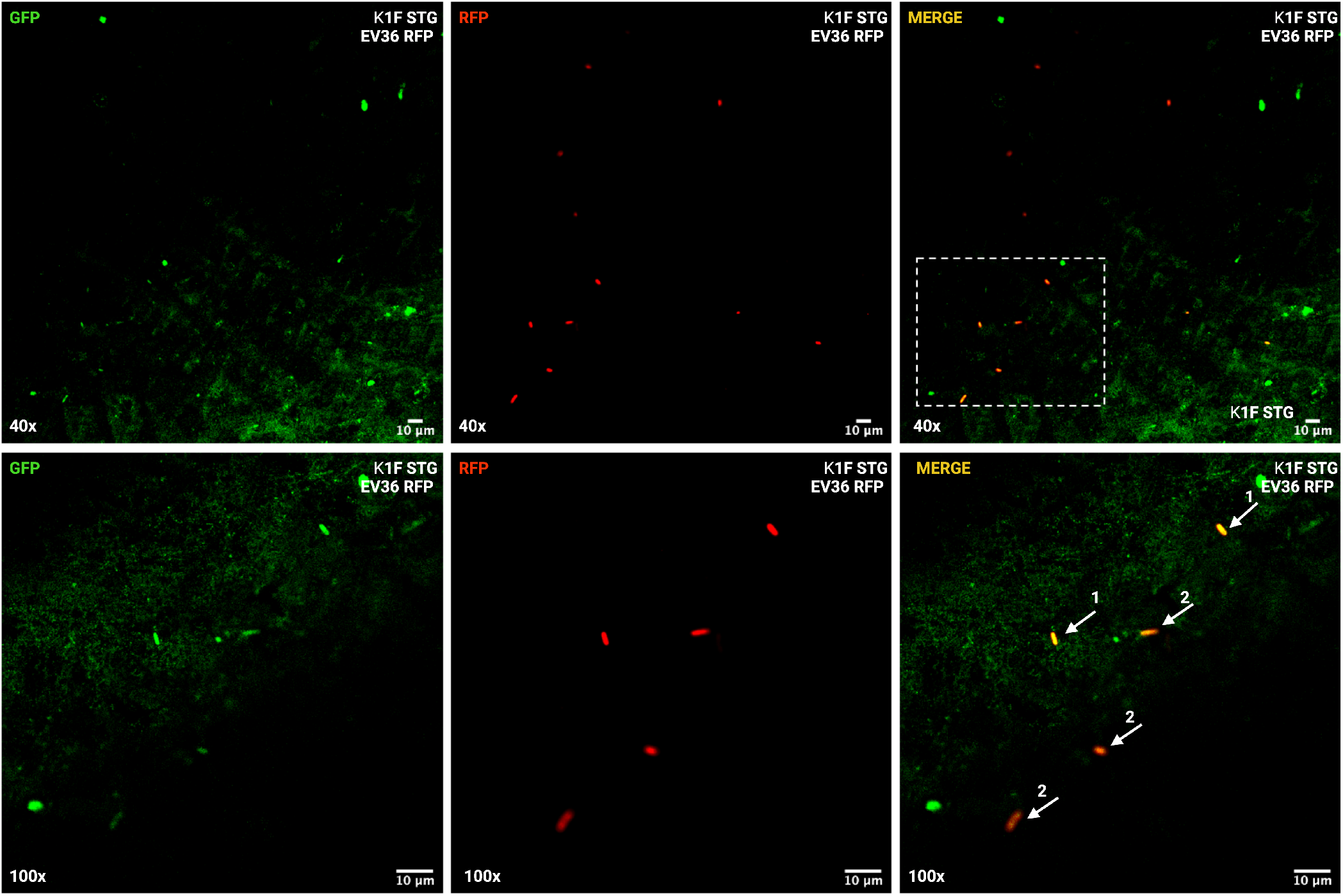
Bacterial capture by directionally immobilised K1F-STG on PDMS. Representative fluorescence microscopy images of K1F STG immobilised on the surface of PDMS devices capturing their host EV36 expressing RFP. When imaging at 40x magnification **(Top)**, immobilised K1F-STG is visible as punctate green dots. Areas without K1F-STG phage immobilisation show free-floating bacteria which appear red in the merge panel towards the top of the field of view. The bottom panel is a 100x magnification of the area marked by the dotted line in the 40x merge panel. In this section, it is possible to visualise captured and infected bacteria **(1)** as well as infected and motile bacteria **(2)**. Captured and infected bacteria appear yellow/orange in the merge panel as the RFP expressing EV36 bacteria are now also expressing GFP due to K1F-STG replication. These bacteria are in focus as they are stationary due to being captured by the immobilised phage. In contrast, infected motile bacteria while also appearing yellow/orange in the merge panel, are not in focus and are moving through the field of view.

The findings presented thus far highlight the need for careful consideration of protein design when engineering biosensor devices. The change in the fusion protein orientation of SpyCatcher to BslA maximised the functional utility of SpyCatcher proteins immobilised on PDMS surface for covalent coupling to SpyTag phage. These experiments thus confirm the robustness of the method of phage immobilisation on PDMS surfaces. Detailed methods on protein expression and device fabrication is outlined in the Supplementary Materials.

### Bacterial capture by K1F-STG Immobilised on PDMS Surface

Following the confirmation of K1F STG on the surface of BslA-SpyCatcher decorated PDMS chips through fluorescence microscopy, we next sought to investigate the viability of the immobilised phage in capturing its cognate host. For these experiments, K1F-STG was immobilised on BslA-SpyCatcher decorated PDMS chips as previously described. The phage immobilised devices were subsequently stored in SM buffer for 3 weeks before performing bacterial capture experiments, to investigate the long-term viability of immobilised phage. For bacterial capture, a 3 day old culture of *E. coli* EV36 bacteria expressing RFP was added on top of the device and incubated for 8 minutes. It was necessary to use a stationary phase culture and a short incubation time to minimise the likelihood of phage-mediated lysis of the host before the acquisition of a fluorescence microscopic image of phage-captured bacteria. Following the incubation of the stationary phase culture incubation, the bacteria were washed off with water using a spray bottle. The rinsed devices were blotted on sterile paper towels to dry. Examination of the PDMS devices under fluorescence microscopy revealed clear evidence of bacterial capture by immobilised K1F-STG by way of co-localised GFP and RFP fluorescent signals. The surface of the device contained three different bacterial populations. Free-floating (uncaptured) and uninfected bacteria were visible as motile red bacteria in the RFP channel only. Motile bacteria appeared out of focus in the acquired images. Bacteria captured and infected by phage appeared with the classical *E*.*coli* rod shape and were in focus and stationary in the field of view. Captured bacteria appeared in both the RFP channel and the GFP channel due to the replication of K1F-STG phage within these cells. Finally, the final population of bacteria also appeared in both the RFP and GFP channels but were motile. These bacteria give GFP fluorescence as a result of infection by free-floating K1F-STG produced from a lysis event from a captured bacteria elsewhere on the device. Taken together, these results clearly demonstrate the ability of K1F-STG covalently immobilised on the surface of PDMS chips to capture their host *E. coli* EV36 and serve as a proof of concept for the use of directionally immobilised phage within a bacterial detection microfluidic device. Further microscopy data, including control devices incubated with K1F-WT, K1F-GFP and bacterial capture on SpyCatcher-BslA devices, are provided in Supplementary Figures S6 and S7.

### Conclusions and Future Directions

The results presented in this manuscript demonstrate a key proof of concept milestone in the creation of a phage-based point of care diagnostic device for bacterial detection. Building on our previous work on the creation of the SpyPhage platform for rapid and modular phage engineering, a simple strategy was devised for the immobilisation of Spy-Tagged phage onto PDMS-based microfluidic devices. The strategy for SpyCatcher fusion protein trapping on PDMS is simple and does not require complex pre-treatment processes. The physical immobilisation of SpyCatcher is achieved by simply spotting and drying as little as 1-5 µl of SpyCatcher fusion protein solution in carbonate buffer onto a hydrophobically treated device mould and pouring over PDMS. Once cured, the PDMS layer is peeled off from the mould resulting in the surface entrapment and display of SpyCatcher proteins on the underside of the PDMS. The SpyCatcher displaying PDMS chip is then ready to be incubated with SpyTag modified phage, resulting in directionally immobilised phage.

During method development, various SpyCatcher fusions were tested for optimised surface display of SpyCatcher active sites. Through rational protein design and rapid prototyping, the fusion of SpyCatcher to the hydrophilic terminus of the bacterial hydrophobin BslA was identified as the optimal spotting solution protein. Devices created using the BslA-SpyCatcher fusion dramatically increased the immobilisation efficiency of SpyTagged phage as visualised by fluorescence microscopy. Despite harsh post-production wash steps and sonication, the phage remained directionally immobilised on the surface of the PDMS device by way of the covalent linkage between SpyTag and SpyCatcher. Furthermore, PDMS chips with surface immobilised phage were able to capture and infect EV36 *E*.*coli* cells thus proving the retention of lytic activity of immobilised phage for up to 3 weeks when devices are stored in SM buffer.

To better envision the utilisation of this technology within a diagnostic/susceptibility testing device, let us consider the general design of a P.O.C microfluidic device as illustrated in Figure 9. The design consists of a (1) the “inlet hole” where the sample to be tested enters the device, (2) the “single-cell phage infection zone” where nanoluciferase signal encoded SpyTag Phage are immobilised and ready to infect bacteria in the sample, (3) the “lysis + Magnetic Nanoparticle (MNP) coupling chamber” where infected cells are lysed and the signal couples to the MNPs, and (4) the “imaging chamber” where the MNPs carrying signal are isolated from sample debris and combined with the substrate for nanoluciferase thus producing a bio-luminescent signal. An imaging device (smartphone/photomultiplier) can then be used to detect the bio-luminescence signal and give a readout.(*22*)

**Figure 9:**
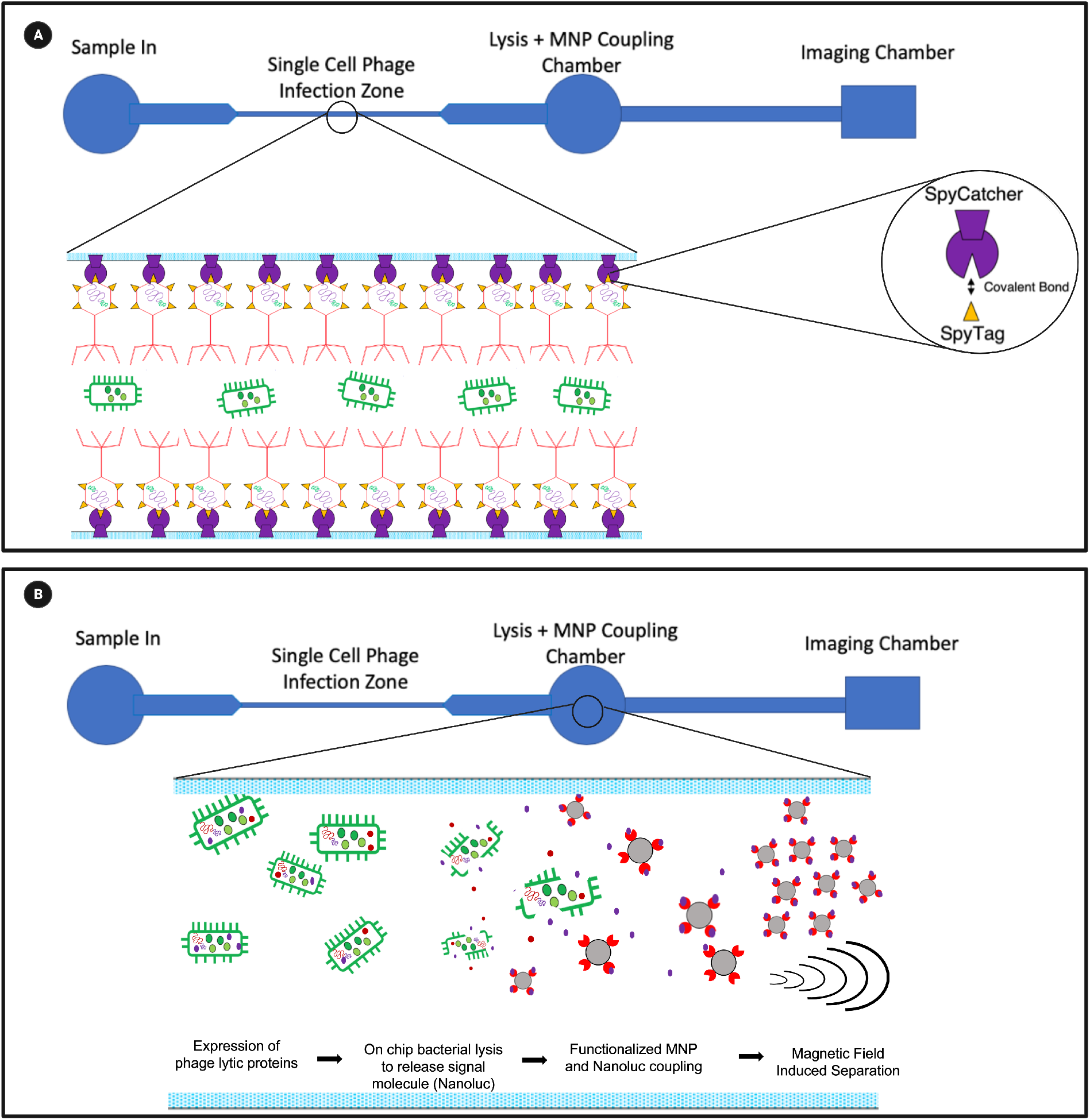
Design of Point of Care Phage Diagnostic Device. The POC device design consists of (1) an inlet whole where the sample to be tested enters the device, (2) single-cell phage infection zone where engineered phage are immobilised and ready to infect bacteria in the sample, (3) Lysis + MNP coupling chamber where infected cells are lysed and signal couples to MNP, and (4) Imaging chamber where MNPs carrying signal are isolated from sample debris and an imaging device gives a readout. SpyTag phage are directionally immobilised into the single-cell phage infection zone of the PDMS-based device using the SpyTag/Catcher system **(A)**. During infection, SpyTag phage infect bacteria with Nanoluc protein pre-packaged into the capsid head. The Nanoluc protein is released from the bacteria during phage-induced lysis and is separated using functionalised magnetic nanoparticles (MNPs) **(B)**.

The use of a microfluidic platform for bacterial detection is advantageous for several reasons. Firstly, microfluidic devices enable precise control of fluid flow and thus spatiotemporal control over reaction conditions and the timing of their occurrence.(*23*) With respect to the concept proposed, the sample to be tested must undergo several stages of processing. Phage infection, bacterial lysis, coupling of signals to magnetic nanoparticles (MNPs) and the reporter signal assay must all occur sequentially and separately from each other.

The use of microfluidics can also help to increase the sensitivity of the bacterial detection system due to their capacity for highresolution separation.(*24*) In order to maximise the chance of infection of target bacteria by signal-carrying phage, the infection zone of the device is designed such that it only allows the passage of single bacterial cells (*24, 25*) . This reduction in reaction space enables the rapid characterisation of a heterogeneous sample by increasing the likelihood of phage infection.(*26*) Following infection of bacteria by phage, released signal molecules are captured by magnetic nanoparticles. The movement of signal-loaded MNPs in the device is facilitated by the use of a handheld magnet, providing a strategy for simple signal amplification and isolation. This ensures that the signal molecules are separated from all cell debris and other background material thereby increasing the signal-to-noise ratio and by proxy; the sensitivity of detection. Furthermore, if the immobilised signal carrying phage can be engineered to be replication-deficient, infection events will not result in the production of virulent phage particles and thereby further reduce the signal-to-noise ratio. In such an event, the replication-deficient or sterile phage could be engineered to only produce lysis proteins and the signal protein which would be expressed by the infected host. Such a design consideration would serve to overcome regulatory hurdles for a phage based point of care device.

This versatile strategy can enable the coupling of different SpyTagged phage to different parts of a microfluidic device by selective and sequential surface exposure. In doing so, a single device can be easily fabricated to detect multiple bacterial pathogens through the facile directional immobilisation of different bacteriophage in an array format. To this end, the methods described herein allow for the conceptualisation of phage based P.O.C devices where the initial identification of the pathogen not only serves to uncover the cause of infection but also assesses the susceptibility of the infecting bacterium to an array of immobilised phage. This dual-purpose approach provides actionable data for immediate therapeutic intervention, enabling the tailored use of phage therapies right from the onset of an infection. Such technology could revolutionize clinical practices by allowing for the customization of treatments based on the specific bacterial strains identified in patients.(*27*) This not only enhances treatment outcomes but also plays a critical role in combating the rising challenge of antibiotic resistance. By integrating diagnostic and therapeutic capabilities, this model ensures that phage therapies are appropriately matched to the pathogen involved, optimizing the efficacy of the intervention and reducing the time to treatment. Through a combination of rational protein design and an orthogonal method for the directional immobolisation of of phage on to PDMS, we hope this work provides generally applicable solutions to increase sensitivity and reduce the time of detection for the next generation of phage-based biosensors.

## 3 Supporting Information

List of plasmids used, List of strains used, donor cassettes for homologous recombination, plasmid architectures for expression constructs, SDS PAGE gels of over expressed SpyCatcher fusions to BslA, SDS PAGE gels of SpyCather-SpyTag coupling reactions, fluorescence microscopy of K1F STG immobilised on SpyCatcher-BslA devices and on-chip capture of bacteria, Negative control fluorescence microscopy experiments using K1F-GFP and K1F-WT, detailed methods for protein expression, device fabrication, on-chip bacterial capture experiments, additional Figures S1-S7 and Tables S1-S3.

## 4 Author Information

### Author Contributions

S.B.W.L. conceived the project, executed experiments and prepared manuscript. J.W. performed fluorescence microscopy imaging. S.B.W.L. designed plasmids for the creation of phage K1F-STG. A.Y.B. and T.F. created phage K1F-STG. Phage K1F-WT, phage K1F-GFP, host strain *E. coli* EV36, and reagents for microscopy were provided by A.P.S. Manuscript was revised by J.P.W., T.F. and V.K. Conflicts of Interest: None

## Acknowledgments

We thank Dr Ella Lucille Thornton (University of Edinburgh) for providing the pE28-BslA-SpyCatcher plasmid and advising on BslA protein expression and handling. This research is supported by EPSRC [UK Research and Innovation (UKRI) Gateway to Research Project Reference: 1917070], by award ACC101854, by the Medical and Life Science Research Fund Bursary [70442] and by the EPSRC/BBSRC grant BB/M017982/1 to the Warwick Integrative Synthetic Biology Centre. T.F. was supported by the Biotechnology National Laboratory of Hungary. J.W. was supported by UKRI BBSRC [Project reference 2266975]. A.P.S. was supported by BBSRC Future Leader’s Fellowship [Grant Reference: BB/N011872/]. S.B.W.L. and V.K.. provided funds for this project. V.K. is a Tata Consultancy Services Affiliate Faculty.

This is an invited contribution from the 15th International Workshop on Bio-Design Automation (IWBDA). A preliminary 2-page version of this manuscript was presented at IWBDA 2021.

## Notes

### Competing Interest Statement

The authors have declared no competing interest.

## References

1. Farooq, U., Yang, Q., Ullah, M. W., and Wang, S. (2018) Bacterial biosensing: Recent advances in phage-based bioassays and biosensors. Biosensors and Bioelectronics 118, 204–216.

2. Anany, H., Brovko, L., El Dougdoug, N. K., Sohar, J., Fenn, H., Alasiri, N., Jabrane, T., Mangin, P., Monsur Ali, M., Kannan, B., Filipe, C. D. M., and Griffiths, M. W. (2018) Print to detect: a rapid and ultrasensitive phage-based dipstick assay for food-borne pathogens. Analytical and Bioanalytical Chemistry 410, 1217–1230.

3. Tolba, M., Minikh, O., Brovko, L. Y., Evoy, S., and Griffiths, M. W. (2010) Oriented immobilization of bacteriophages for biosensor applications. Applied Environmental Microbiology 76, 528–35.

4. Burnham, S., Hu, J., Anany, H., Brovko, L., Deiss, F., Derda, R., and Griffiths, M. W. (2014) Towards rapid on-site phage-mediated detection of generic Escherichia coli in water using luminescent and visual readout. Analytical and Bioanalytical Chemistry 406, 5685–93.

5. Chen, J., Alcaine, S. D., Jiang, Z., Rotello, V. M., and Nugen, S. R. (2015) Detection of Escherichia coli in drinking water using T7 bacteriophage-conjugated magnetic probe. Analytical Chemistry 87, 8977–84.

6. Huang, C., Mahboubat, B. Y., Ding, Y., Yang, Q., Wang, J., Zhou, M., and Wang, X. (2021) Development of a rapid Salmonella detection method via phage-conjugated magnetic bead separation coupled with real-time PCR quantification. LWT 142.

7. Laube, T., Cortes, P., Llagostera, M., Alegret, S., and Pividori, M. I. (2014) Phagomagnetic immunoassay for the rapid detection of Salmonella. Applied Microbiology and Biotechnology 98, 1795–805.

8. Shabani, A., Marquette, C. A., Mandeville, R., and Lawrence, M. F. (2013) Magnetically-assisted impedimetric detection of bacteria using phage-modified carbon microarrays. Talanta 116, 1047–53.

9. Smartt, A. E., and Ripp, S. (2011) Bacteriophage reporter technology for sensing and detecting microbial targets. Analytical and Bioanalytical Chemistry 400, 991– 1007.

10. Wang, C., Sauvageau, D., and Elias, A. (2016) Immobilization of Active Bacteriophages on Polyhydroxyalkanoate Surfaces. ACS Applied Material Interfaces 8, 1128– 38.

11. Rosner, D., and Clark, J. (2021) Formulations for Bacteriophage Therapy and the Potential Uses of Immobilization. Pharmaceuticals (Basel) 14.

12. Gordon, S. M., Srinivasan, L., and Harris, M. C. (2017) Neonatal Meningitis: Overcoming Challenges in Diagnosis, Prognosis, and Treatment with Omics. Frontiers in Pediatrics 5, 139.

13. King, J. E., Aal Owaif, H. A., Jia, J., and Roberts, I. S. (2015) Phenotypic Heterogeneity in Expression of the K1 Polysaccharide Capsule of Uropathogenic Escherichia coli and Downregulation of the Capsule Genes during Growth in Urine. Infection and Immunity 83, 2605–13.

14. Krishnan, S., Chang, A. C., Stoltz, B. M., and Prasadarao, N. V. (2016) Escherichia coli K1 Modulates Peroxisome Proliferator-Activated Receptor gamma and Glucose Transporter 1 at the Blood-Brain Barrier in Neonatal Meningitis. Journal of Infectious Diseases 214, 1092–104.

15. Moller-Olsen, C., Ho, S. F. S., Shukla, R. D., Feher, T., and Sagona, A. P. (2018) Engineered K1F bacteriophages kill intracellular Escherichia coli K1 in human epithelial cells. Scientific Reports 8, 17559.

16. Zakeri, B., Fierer, J. O., Celik, E., Chittock, E. C., Schwarz-Linek, U., Moy, V. T., and Howarth, M. (2012) Peptide tag forming a rapid covalent bond to a protein, through engineering a bacterial adhesin. Proceedings of the National Academy of Sciences 109, E690–7.

17. Liyanagedera, S., Williams, J., Wheatley, J., Biketova, A., Hasan, M., Sagona, A., Purdy, K., Puxty, R., Feher, T., and Kulkarni, V. (2022) SpyPhage: A Cell-Free TXTL Platform for Rapid Engineering of Targeted Phage Therapies. ACS Synthetic Biology 11.

18. Thornton, E. L., Paterson, S. M., Gidden, Z., Horrocks, M. H., Laohakunakorn, N., and Regan, L. (2022) Self-Assembling Protein Surfaces for In Situ Capture of Cell-Free-Synthesized Proteins. Frontiers in Bioengineering and Biotechnology 10.

19. Heyries, K. A., Marquette, C. A., and Blum, L. J. (2007) Straightforward Protein Immobilization on Sylgard 184 PDMS Microarray Surface. Langmuir 23, 4523– 4527.

20. Liu, W., Li, S., Wang, Z., Yan, E. C. Y., and Leblanc, R. M. (2017) Characterization of Surface-Active Biofilm Protein BslA in Self-Assembling Langmuir Monolayer at the Air-Water Interface. Langmuir 33, 7548–7555.

21. Park, S., Liu, L., Demirkır van der Heijden, O., Lohse, D., Krug, D., and Koper, M. T. M. (2023) Solutal Marangoni effect determines bubble dynamics during electrocatalytic hydrogen evolution. Nature Chemistry 15.

22. Alonzo, L. F., Jain, P., Hinkley, T., Clute-Reinig, N., Garing, S., Spencer, E., Dinh, V. T. T., Bell, D., Nugen, S., Nichols, K. P., and Le Ny, A.-L. M. (2022) Rapid, sensitive, and low-cost detection of Escherichia coli bacteria in contaminated water samples using a phage-based assay. Scientific Reports 12, 7741.

23. Benedetto, A., Accetta, G., Fujita, Y., and Charras, G. (2014) Spatiotemporal control of gene expression using microfluidics. Lab on a Chip 14, 1336–47.

24. Hong, S. C., Kang, J. S., Lee, J. E., Kim, S. S., and Jung, J. H. (2015) Continuous aerosol size separator using inertial microfluidics and its application to airborne bacteria and viruses. Lab on a Chip 15, 1889–97.

25. Lu, H., Caen, O., Vrignon, J., Zonta, E., El Harrak, Z., Nizard, P., Baret, J. C., and Taly, V. (2017) High throughput single cell counting in droplet-based microfluidics. Scientific Reports 7, 1366.

26. Zhang, D., Bi, H., Liu, B., and Qiao, L. (2018) Detection of Pathogenic Microorganisms by Microfluidics Based Analytical Methods. Analytical Chemistry 90, 5512– 5520.

27. Wheatley, J. P., Liyanagedera, S. B. W., Fehér, T., Sagona, A. P., and Kulkarni, V. (2024) TXTL-Powered K1F Internal Capsid Protein Engineering for Specific, Orthogonal and Rapid Phage-based Pathogen Detection. Biorixv

