## Supporting Information for "Directional Immobilisation of SpyTag Bacteriophage on PDMS surfaces for Phage based Microfluidics"

Contents:


| Item | Title | Page |
|---|---|---|
| Table S1 | List of plasmids | 2 |
| Table S2 | List of strains | 2 |
| Table S3 | pUC_GFP-ST7 donor cassette sequence for creation of K1F-STG | 2 |
| Figure S1 | Gene10b::GFP-SpyTag synthetic gene cassette | 3 |
| Figure S2 | Plasmid architectures of expression constructs | 4 |
| Figure S3 | SpyCatcher-Bsla and Bsla-SpyCatcher over expression | 5 |
| Figure S4 | SpyCatcher-SpyTag coupling reaction | 5 |
| Figure S5 | Fluorescence microscopy images of K1F-STG directionally immobilised on SpyCatcher-BslA surface-displayed PDMS device. | 6 |
| Figure S6 | Fluorescence microscopy images of bacterial capture by K1F-STG directionally immobilised on SpyCatcher-BslA surface-displayed PDMS device. | 7 |
| Figure S7 | Fluorescence microscopy images of negative control devices with K1F-GFP | 8 |

**Table S1.** List of plasmids.

| Name of plasmid | Construction/Source | Purpose |
|---|---|---|
| **p70a-GFP-SpyTag** | Gibson assembly of SpyTag synthetic gene cassette into pBEST-p70b-UTR1-deGFP-6xHis-T500 (Addgene # 92222) | Covalent coupling to PDMS surface displayed SpyCatcher |
| **p70a-SpyCatcher-mCherry** | SpyCatcher-mCherry synthetic gene cassette cloned into pBEST-p70b-UTR1-deGFP-6xHis-T500 (Addgene # 92222) to replace deGFP | Creation of PDMS chips with surface displayed SpyCatcher |
| **pET28-GST-BslA-SpyCatcher** | Kind gift from Dr Ella Thornton (University of Edinburgh) | Creation of PDMS chips with surface displayed SpyCatcher |
| **T7p14-GST-SpyCatcher-BslA** | Three-piece Gibson assembly of SpyCatcher and BslA into T7p14 backbone derived from T7p14-GFP (Arbor #502134) | Creation of PDMS chips with surface displayed SpyCatcher |

**Table S2.** List of strains.

| Strain | Origin | Purpose |
|---|---|---|
| **EV36-RFP** | Kind gift from Dr Antonia Sagona (University of Warwick) | Host strain of K1F phage expressing RFP |
| **BL21 (DE3)** | Thermo Fisher (EC0114) | Expression of SpyCatcher-BslA and BslA-SpyCatcher |
| **NEB Turbo** | New England Biolabs (C2984I) | Expression of sigma 70 driven expression of GFP-SpyTag and SpyCatcher-mCherry |

| **pUC_GFP-ST7 donor cassette sequence** |
|---|
| GGTGATGGTCCTGTTCTGCTGCCAGACAATCACTATCTGAGCACGCAAAGCGTTCTGTCTAAAG<br>ATCCGAACGAGAAACGCGATCATATGGTTCTGCTGGAGTTCGTAACCGCAGCGGGCATCACGCA<br>TGGTATGGATGAACTGTACAAAGGTAGTGGTGAAAGTGGTGAAAACCTGTATTTTCAGGGAGCC<br>CACATCGTGATGGTGGACGCCTACAAGCCGACGAAGTAGTAGTAATTGAAACCCCTTGGGTGCC<br>TTCGGGTGCTTGAGGGGTTTTTGCTTAAAGTGAGAGGAGACTTATGGCTCAATACATTCCACTG<br>AATGCTAACGATGACTTAGATGCCATCAACGATATGTTAGCTGCTATCGGTGAACCAGCAGTCC |

**Table S3.** pUC_GFP-ST7 donor cassette sequence for creation of K1F-STG

DNA sequence of the donor cassette (excluding cut sites) for cloning into pUC19 vector to create donor plasmid pUC_GFP-ST7. Sequence contains 5’ and 3’ homology arms for GFP and gp11, respectively. Sequence design enables SpyTag gene fusion to GFP of gene10b::GFP in K1F-GFP to create K1F-STG *via* homologous recombination.

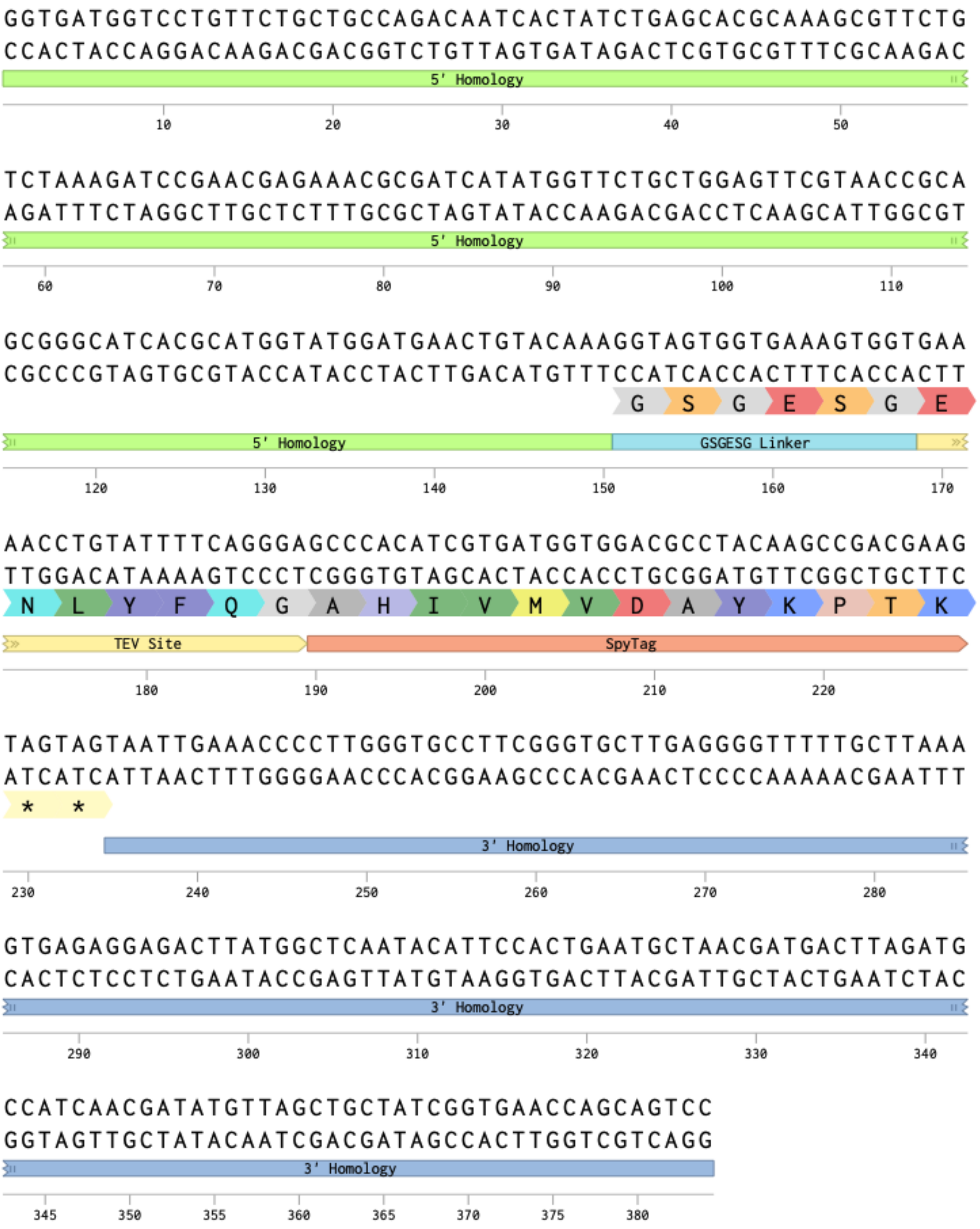


**Figure S1.** gene10b::GFP-SpyTag synthetic gene cassette annotated map

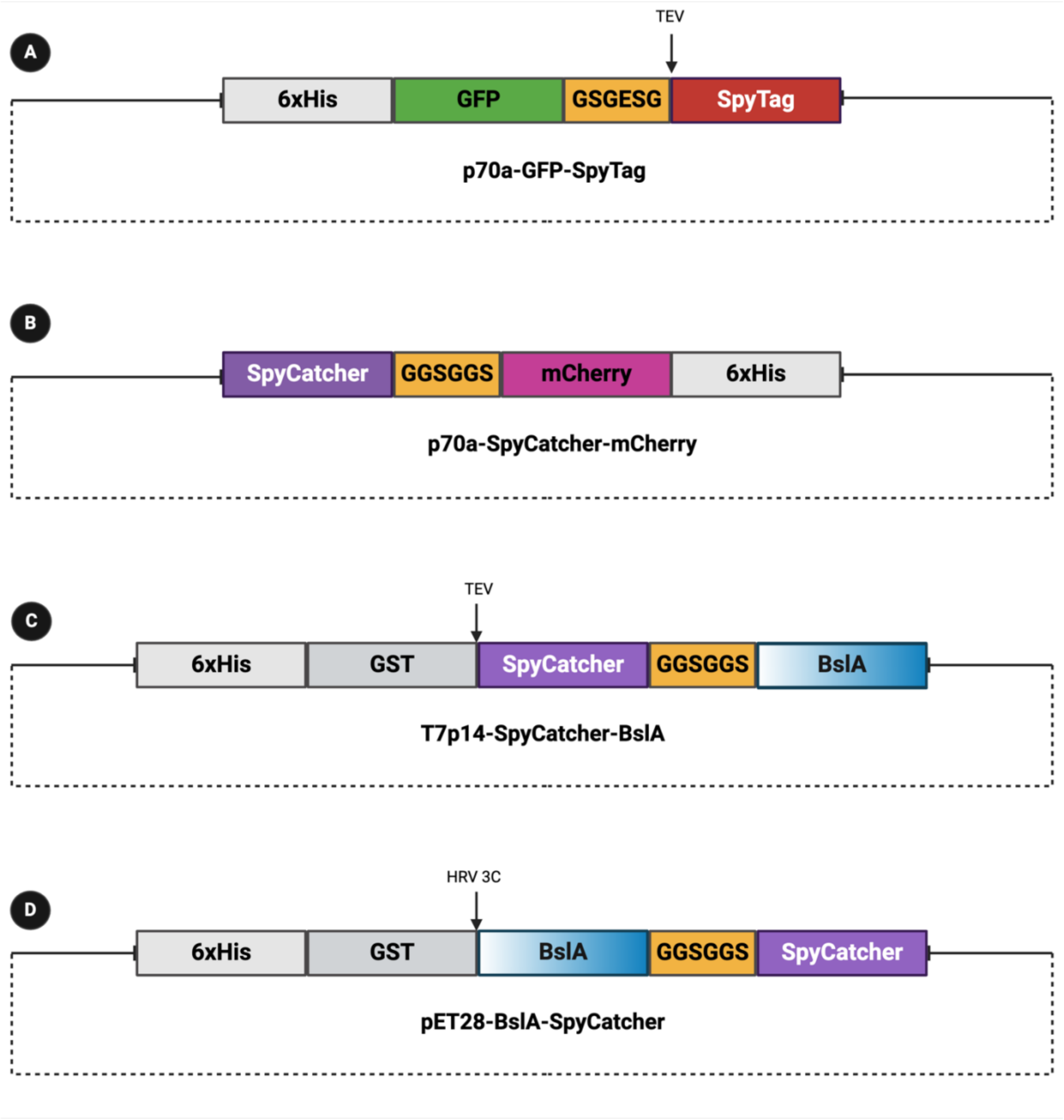


**Figure S2.** Plasmid architectures of expression constructs used in this study.

GFP fusion to SpyTag was created to allow for simple testing of Immobilisation strategy prior to attempting to immobilise SpyTag phage. TEV cleavage site was incorporated down stream of the linker to replicate the design of the SpyTag fusion to minor capsid of K1F (A). SpyCatcher fusion to mCherry was used during method development of immobilisation strategy, however this fusion did not lead to surface displayed SpyCatcher (B). SpyCatcher fusion to hydrophobic N-terminus of BslA with a TEV cleavage site for optional removal of GST solubility tag and 6x His tag (C). SpyCatcher fusion to hydrophilic terminus of BslA with a HRV 3C cleavage site for optional removal of GST solubility tag and 6x His tag (D). Gradient of BslA block indicates the Hydrophobic (white) and hydrophilic (blue) regions of the protein.

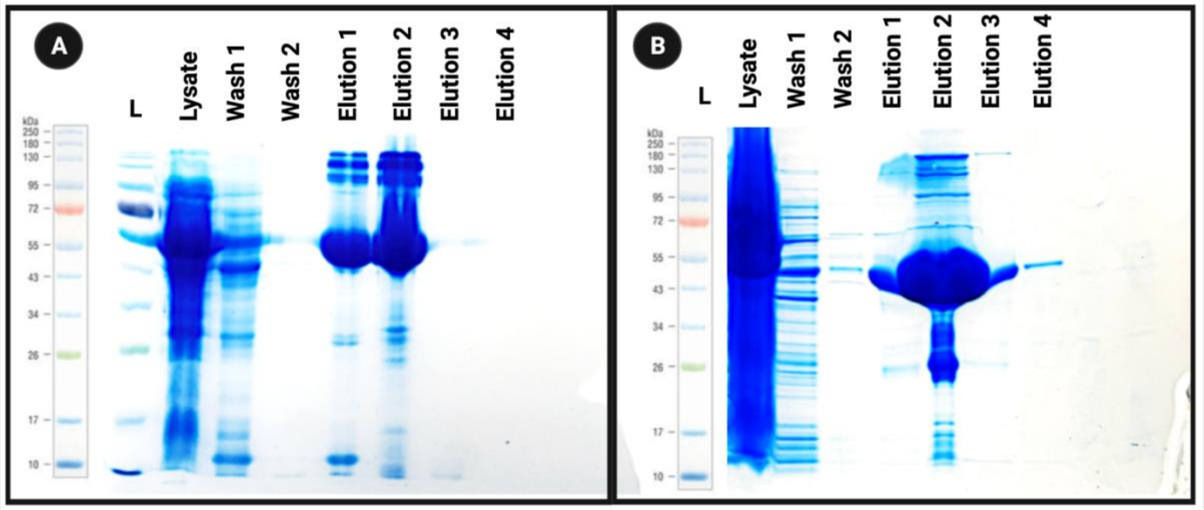


**Figure S3.** SpyCatcher-BslA and BslA-SpyCatcher over expression.

Initial expression trials revealed the accumulation of BslA fusions to SpyCatcher in the insoluble fraction of expressed cultures. As such a rapid expression protocol in which cultures were grown at 40°C for 5 hours at 250 RPM was developed. This led to dramatic improvements in the expression of both SpyCatcher-BslA (A) and BslA-SpyCatcher (B).

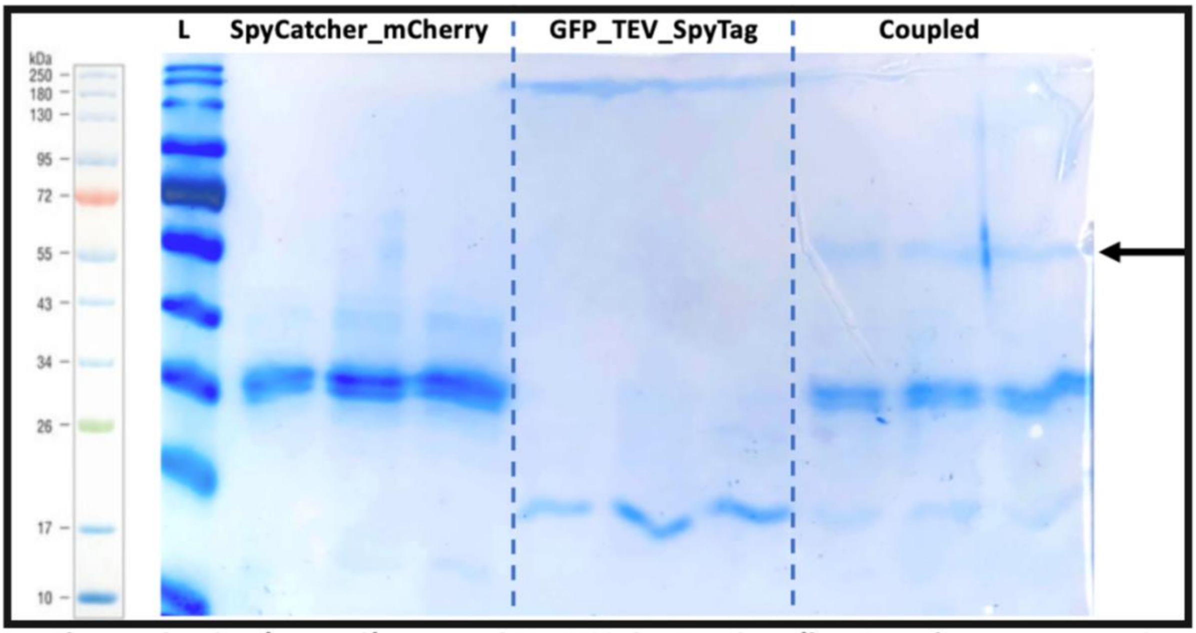


**Figure S4.** SpyCatcher- SpyTag Coupling Reaction

Coupling reaction of SpyCatcher-mCherry and GFP-SpyTag results in the formation of a covalent complex which has the cumulative mass of both proteins (black arrow). This demonstrates the ability of SpyCatcher to form a covalent link with SpyTag which can withstand the conditions of a denaturing PAGE gel.

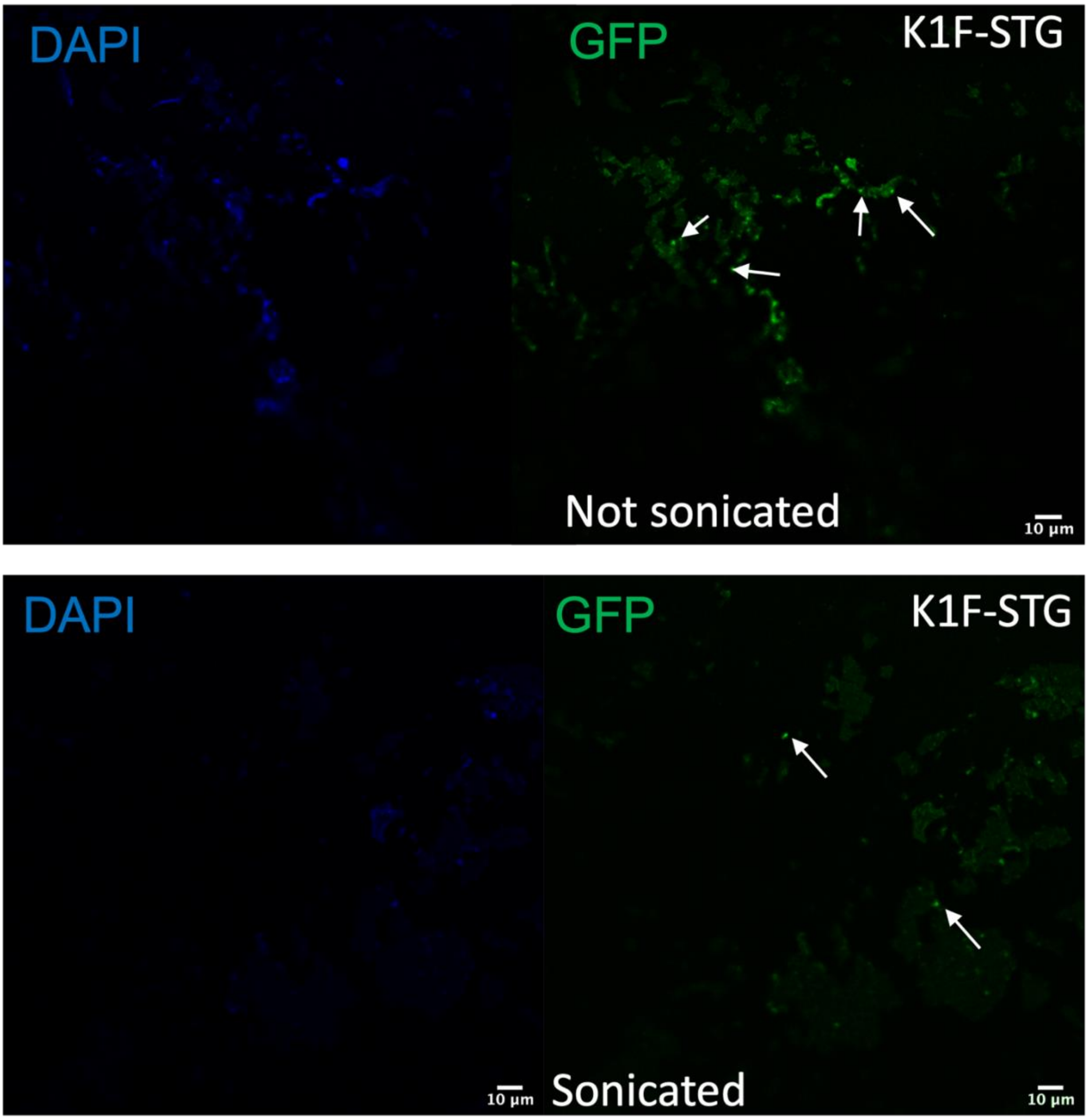


**Figure S5.** Immobilisation of K1F-STG on PDMS devices displaying SpyCatcher-BslA.

Fluorescence microscopy image of the surface of PDMS device before (Top) and after (Bottom) sonication. Phage are visible as punctate green fluorescent dots on the surface of the device even after 30 minutes of surface disruption in a sonicating water bath. The overlap of the green and blue fluorescent channels indicate the presence of phage due to the fluorescence from GFP on the capsid head and the DAPI stained phage DNA respectively. The results indicate the strength of the covalent interaction between the SpyTag on the capsid of the phage and the SpyCatcher-BslA fusion immobilised on the PDMS surface.

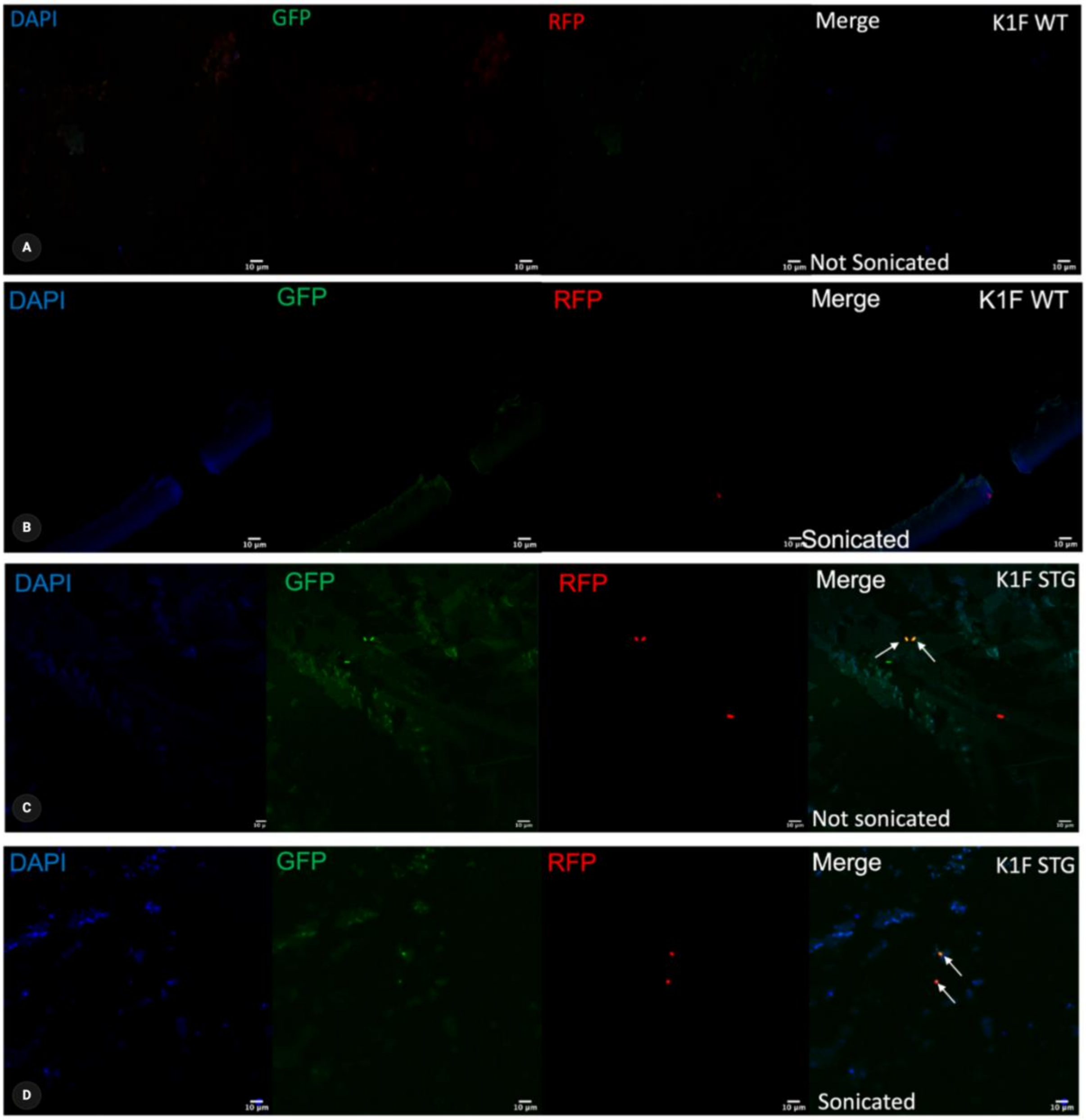


**Figure S6.** Bacterial capture by K1F-STG immobilised via SpyCatcher-BslA on PDMS

Devices made with SpyCatcher-BslA proteins could be used for the directional immobilisation of K1F-STG phage. These phage immobilised devices were able to capture bacteria on-chip, though at a lower efficiency compared to BslA-SpyCatcher devices (C and D). Control SpyCatcher-BslA devices were incubated with K1F-WT phage which lacks SpyTag (A and B). Staining with DAPI did not reveal the capsid structures of phage, thus demonstrating that SpyTag on the capsid is required for directional immobilisation on SpyCatcher decorated devices. These control devices were also unable to capture bacteria on-chip. White arrows indicate captured bacteria.

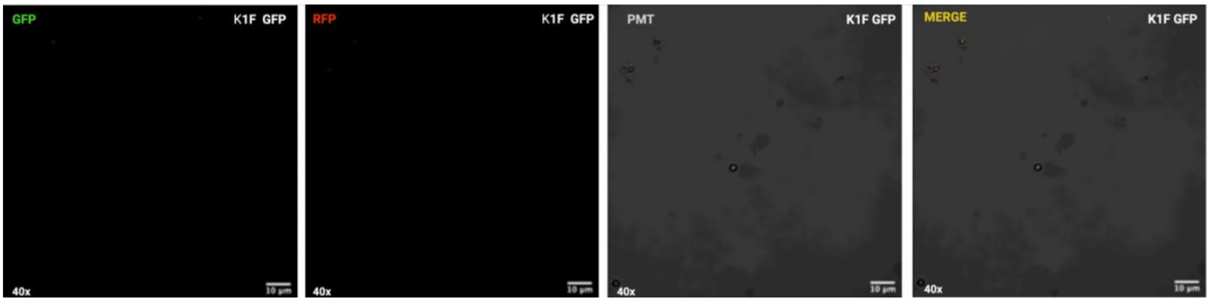


**Figure S7.** K1F-GFP negative control devices.

Devices made with BslA-SpyCatcher or SpyCatcher-BslA proteins are unable to be used for the directional immobilisation of K1F-GFP phage due to the lack of SpyTag protein on the capsid of K1F-GFP. Repeated microscopic analysis did not reveal any K1F-GFP phage immobilised on the surface of the device thus demonstrating the requirement of SpyTag for directional immobilisation with surface exposed SpyCatcher. Such devices were also unable to capture bacteria. The use of the Photomultiplier (PMT) setting only revealed background fluorescence from the device.

### S1. Bacteriophage Methods

Routine phage methods i.e. phage propagation from liquid *E. coli* cultures, titering, plaque assay and plaque PCR were carried out as described previously, unless stated otherwise[1]. Propagation of K1F variants was performed as described in Supporting Information S5. PEG precipitated phage stocks were stored in sterile 15 mL falcon tubes at 4°C until use and plaque assayed to determine Plaque Forming Units (PFU).

### S2. Homologous Recombination and CRISPR/Cas9 Selection

K1F-gene10b::GFP-SpyTag (K1F-STG) phage was engineered using plasmid based homologous recombination, facilitated by CRISPR/Cas9 selection, as described previously[1].

Briefly, K1F-STG was engineered by growing the K1F-GFP phage (carrying g10b::GFP) *on E. coli* EV36/pUC-GFP-ST. The resulting phage mix was enriched on *E. coli* EV36 + pCas9GFP-ST for three rounds, followed by a plaque assay and plaque PCR to screen for engineered phage. The engineered segments of the phage were PCR-amplified for verification by Sanger sequencing. Donor cassette sequence and design of pUC-GFP-ST is provided in Supporting Information Table S3 and Figure S1 respectively.

### S3. Protein Expression

SpyCatcher-mCherry and GFP-SpyTag were expressed in NEB Turbo cell lines. Both proteins were expressed in 1L cultures in 2.5L baffled flasks and incubated at 25°C for 8 hours without IPTG induction before harvesting. SpyCatcher-BslA and BslA-SpyCatcher were expressed in BL21(DE3) cell lines. Both proteins were expressed in 1L cultures in 2.5L baffled flasks and incubated at 37°C until OD600 of 0.5 was reached at which point cultures were induced with 1 mM IPTG and grown for a further 5 hours at 40°C before harvesting. Following protein expression, cell cultures were pelleted at 7000 RPM for 10 min at 4°C. Cell pellets were resuspended in Lysis Buffer (4°C) at a ratio of 50 mL buffer per litre of culture. Following resuspension, cells were lysed using sonication using the Sonics Vibra-Cell VCX130 ultrasonic processor (settings: amplitude = 80%, pulse on = 10 s, pulse off = 30 s, time = 3 min). Cell lysates were centrifuged at 30 000 g for 60 min at 4°C to separate the soluble and insoluble fractions. The soluble fraction was decanted out and stored on ice, as were the insoluble fractions. Prior to His affinity purification of the soluble fraction, the sample was passed through a 0.22 µM filter. All passages through the His Trap columns were done using a 5 ml sterile syringe. First, the column was washed with dH20 to get rid of ethanol present in the column from the man-ufacturer/previous user. The column was then equilibrated with 5 ml of resuspension Buffer. 5 ml of filtered soluble cell lysate was passed through the column to bind His tagged proteins of interest. This was repeated until all the soluble cell lysate was loaded into the column. The column was then washed with 15 mL of Lysis buffer, split into two washes of an initial 10 mL and then 5 mL, and collected into two clean 15 mL falcon tubes and put on ice. Finally, an initial elution step was performed with 5 mL of 50 mM Imidazole elution buffer into a clean 15 ml falcon tube. Next, the column was eluted a further 3 times with 3 mL of 300 mM Imidazole per elution and collected into 3 separate clean 15 mL falcon tubes. All washes and elutions collected were stored on ice at 4°C until further analysis via SDS PAGE or downstream use. Protein concentration and purity were determined using Nanodrop. Size exclusion chromatography (SEC) was used to further purify all proteins used in this study. SpyCatcher-mCherry and GFP-SpyTag were stored in protein storage buffer in 2 mg/mL stocks at -20°C. SpyCatcher-BslA and BslA-SpyCatcher were stored in PBS in 2 mg/mL stocks at 4°C (Supplementary Figure S3). Due to the ability of BslA fusions to form monolayers, new pipette tips were used for each liquid handling step of these proteins. All SpyCatcher fusion proteins were buffer exchanged into 0.1 M carbonate buffer (pH = 9) using PD-10 desalting columns immediately prior to their use for device fabrication.

Purified GFP-SpyTag and SpyCatcher-mCherry were coupled in a coupling reaction with equimolar concentrations of each protein for 1 hour. The coupled reaction was analysed by SDS PAGE to demonstrate the covalent linkage between SpyCatcher and SpyTag. The covalent complex appears as a band with a mass of the cumulative weight of both proteins (Supplementary Figure S4).

### S4. PDMS Device Fabrication

All moulds used for device fabrication (silicone wafers, microscope slides, microwell plates or custom moulds) were treated with a general-purpose windscreen rain repellent hydrophobic agent. The moulds were wetted with the hydrophobic agent and allowed to air dry for 10 minutes and excess was removed by blotting on kimtech wipes. Care was taken to treat all surfaces of the mould to ensure thorough hydrophobic treatment. Initial proof of concept work was done by spotting 5-10 µL of desired SpyCatcher fusion onto a hydrophobically treated glass slide or micro-well plate. A similar volume of spotting protein solution was used when creating devices with silicon wafers. To obtain devices with a larger surface area of immobilised SpyCatcher, larger volumes up to 200 µL were spotted on to glass slides or glass coverslip moulds. The spotted solutions were dried onto the mould surface in a 55°C oven. PDMS precursor and curing agent (SYLGARD™ 184 Silicone Elastomer Kit) were mixed in a 10:1 ratio, degassed for 1 hour, and subsequently poured over the spotted mould. Following curing in a 55°C oven, the PDMS chip was cut out and slowly peeled away to create a PDMS chip with surface-decorated SpyCatcher proteins. SpyCatcher decorated PDMS device surfaces were incubated with either GFP-SpyTag in PBS (2 mg/mL) or K1F-STG in PBS (9 Log PFU/mL) for 1 hour to facilitate covalent coupling to the PDMS surface. Following incubation, the GFP-SpyTag/K1F-STG solutions were rinsed from the device using water and the devices were blotted on sterile paper towels. Covalently coupled devices were then placed into a 2L flask containing 1L of water and incubated with agitation at room temperature for 30 minutes. This step was repeated with fresh water for the second round of washing. Following the wash steps, the devices were sonicated in 50 mL falcon tubes containing water in a sonicating water bath set at room temperature for 30 minutes. Devices were subsequently visualised using a protein gel imager or a fluorescence microscope. Phage immobilised devices were stored in SM buffer in 50 mL falcon tubes at 4°C for up to 3 weeks without loss of immobilised phage activity.

### S5. On-Chip Bacterial Capture

For on-chip bacterial capture experiments, phage immobilised devices were removed from their storage buffers and blot dried on sterile paper towels. 100 µL EV36 RFP cultures overgrown from a single colony for 3 days were incubated on top of the phage immobilised devices for 8 minutes to facilitate bacterial capture by phage. Following incubation, the excess bacteria were washed away under a constant stream of water for 30 seconds. The devices were examined immediately under fluorescence microscopy to visualise bacterial capture. As a negative control, devices were fabricated in the same manner with the K1F-GFP phage which lacks SpyTag. In multiple repeats of this control condition, microscopic analysis was unable to locate K1F-GFP non-specifically interacting with the PDMS surface. These devices were also unable to capture bacteria (Supplementary Figure S7). In a similar manner, K1F-WT which also lacks SpyTag was also used as a negative control. These devices did not show phage retention on the surface of the device where phage nucleic acids were stained for by DAPI. These control devices also did not capture bacteria (Supplementary Figure 6).

### S10. Fluorescence microscopy

Fluorescence microscopy was undertaken to visualise the immobilisation of GFP-SpyTag or K1F-STG with SpyCatcher fusions entrapped on PDMS surfaces. These experiments were carried out using the Zeiss LSM 880 laser scanning confocal microscope with Airyscan, with fluorophore excitation occurring at the following wavelengths: DAPI at 405nm, GFP at 488nm and RFP at 561nm. Immobilisation on the surfaces of devices were visualised as co-localisation of DAPI and GFP signals from the phage nucleic acids and GFP capsid modification, respectively on the surfaces of devices. The ability of immobilised phage to capture bacteria were visualised as the co-localisation of GFP and RFP signals from GFP on phage capsids and RFP expressed in host EV36 cells. K1F-WT and K1F-GFP were used as controls as these phages lack SpyTag on their capsid head and are thus unable to covalently bind to SpyCatcher on the surface of PDMS devices.

**Supplementary References**

1. Liyanagedera, S., Williams, J., Wheatley, J., Biketova, A., Hasan, M., Sagona, A., Purdy, K., Puxty, R., Feher, T., and Kulkarni, V. (2022) SpyPhage: A Cell-Free TXTL Platform for Rapid Engineering of Targeted Phage Therapies. ACS Synthetic Biology 11
